# SEASONAL PM_2.5_ DIFFERENTIALLY REGULATES JAK2/STAT3 SIGNALING IN RURAL AND URBAN COHORTS

**DOI:** 10.1101/2025.09.25.678452

**Authors:** Sukanya Ghosh, Rupa Chaudhuri, Meghna Mukherjee, Anurima Samanta, Priyanka Saha, Lucas Henneman, Deepanjan Majumdar, Mita Ray Sengupta, Anindita Chakraborty, Bidisha Maiti, Supratim Ghosh, Avik Biswas, Dona Sinha

## Abstract

Ambient particulate matter (PM_2.5_) is a major environmental carcinogen linked to lung cancer, yet its molecular insights on asymptomatic non-smokers remains unclear. This study examined the effect of seasonal fluctuations of PM_2.5_ on oxidative stress and pro-oncogenic signaling in rural (RU) and urban (UR) cohorts from West Bengal, India. Environmental monitoring revealed higher PM_2.5_ and associated benzo[α]pyrene in UR, during winter, induced oxidative stress (elevated ROS, 8-OHdG), reduced antioxidants (SOD, catalase, GPx), and promoted airway inflammation. Transcriptomic and bioinformatic analyses identified activation of IL-6/EGFR-driven JAK2/STAT3 signaling and its crosstalk with Ras/Raf/MAPK, leading to increased expression of downstream effectors (BCL-2, MCL-1, c-MYC, cyclin D1) and repression of tumor suppressors (BAX, p21). Notably, downregulation of JAK/STAT inhibitors PIAS2 and SOCS2 suggested persistent activation of oncogenic signaling. Linear mixed-effects models linked winter PM_2.5_ surges to oxidative stress, inflammation, and altered JAK2/STAT3 signaling, while regression models showed stronger responses in UR. Risk modeling predicted significantly higher lung cancer mortality in UR, underscoring the role of seasonal PM_2.5_ surges in JAK2/STAT3-driven carcinogenic susceptibility and the urgent need for targeted interventions.

## 1. Introduction

Airborne Particulate Matter (PM_2.5_), with an aerodynamic diameter of ≤2.5 µm, is one of the major air pollutants driving declining air quality and increasing disease burden in densely populated urban areas (Southerland et al. 2022). A recent study projected that in 2021, PM_2.5_ contributed to 0.37 million deaths and 8.9 million Diseases Adjusted Life Years (DALYs) of lung cancer and by 2045 these figures are expected to increase further (Deng et al. 2025). The last decade has seen a sharp rise in PM_2.5_ levels across the Indo-Gangetic Plain, affecting both urban and rural areas (Jaganathan et al. 2025). Nearly 84% of India’s population is exposed to PM_2.5_ concentrations exceeding the annual standards set by the National Ambient Air Quality Standards (NAAQS)-40 µg/m³ and the World Health Organization (WHO)-5 µg/m³ (Ravishankara et al. 2020).

The toxicological effects of PM_2.5_ are attributed to its small size, allowing deep lung penetration, and its complex composition, which includes heavy metals, organic pollutants such as Polycyclic Aromatic Hydrocarbons (PAHs), and Reactive Oxygen Species (ROS) precursors (Loomis et al. 2013; Wang et al. 2025a). Experimental and epidemiological studies have linked PM_2.5_ exposure to oxidative stress, inflammation, DNA damage, epigenetic alterations and dysregulation of signaling pathways implicated in lung carcinogenesis (Chen et al. 2024; Wang et al. 2025b). In particular, activation of pathways such as Janus kinase 2 (JAK2)/Signal transducer and activator of transcription 3 (STAT3) (Hsieh et al. 2022) and Rat Sarcoma Virus (Ras)/Rapidly Accelerated Fibrosarcoma (Raf)/Mitogen-Activated Protein Kinase (MAPK) (Kim et al. 2022) have been associated with PM_2.5_-mediated pro-inflammatory response and oxidative stress. Additionally, crosstalk between MAPK and JAK/STAT pathways have been well evidenced (Liu et al. 2022; Hu et al. 2021). However, epidemiological evidences of dysregulation of the JAK/STAT or the Ras/Raf/MAPK pathways in high PM_2.5_ exposed populations have remained largely undefined.

While the relationship between PM_2.5_ exposure and lung cancer risk has been established in smokers and never smokers, there is limited understanding of early molecular alterations in non-smoking, asymptomatic individuals chronically exposed to rural and urban air pollution. Several studies have established spatial and temporal variation of PM_2.5_ during different seasons (Liu et al. 2019; Zhan et al. 2023; Suthar et al. 2024; Nautiyal et al. 2025). Seasonal fluctuations in PM_2.5_ levels, especially the winter surge common in the Indo-Gangetic plain, may exacerbate the respiratory morbidities (Dutta and Jinsart 2022; Sinha et al. 2024). Reports on the impact of seasonal changes in PM_2.5_ on health effects with underlying molecular alterations are limited. Moreover, rural-urban disparities in exposure and susceptibility remain underexplored, despite differences in pollutant composition, socio-demographics, and lifestyle factors.

This study aimed to investigate the effect of seasonal and spatial variations in ambient PM_2.5_ on oxidative stress response and pro-oncogenic signaling in asymptomatic individuals from urban and rural regions of West Bengal, India. By integrating environmental monitoring, cytological and hematological analyses, transcriptomic profiling, bioinformatic pathway analysis and molecular biological approaches we sought to elucidate the role of oxidative stress and key signaling pathways—including JAK2/STAT3 and Ras/Raf/MAPK—in mediating PM_2.5_-induced pro-carcinogenic alterations. Our findings provide critical insights into the early molecular events associated with chronic PM_2.5_ exposure in never-smoking rural and urban populations of West Bengal, India.

## 2. Results

### 2.1. Spatiotemporal patterns of PM_2.5_ with metal and Benzo[alpha]pyrene (B[α]P) enrichment at RU and UR regions

Monitoring of ambient PM_2.5_ was carried out from Jan-Dec 2023 at Rural (RU) and Urban (UR) sites (Fig. 1a). Jan-Mar, 2023 was considered as winter-spring months and Apr-Oct, 2023 as summer-monsoon months. At the RU air quality monitoring station (Diamond Harbour Women’s University), PM_2.5_ averaged 45.21 ± 12.25 µg/m³ in summer-monsoon and 66.40 ± 20.75 µg/m³ in winter-spring. The RU sampler at Boria showed similar seasonal values (summer-monsoon, 54.99 ± 1.35 vs. winter-spring, 82.54 ± 14.00 µg/m³). At the UR station (University of Calcutta), concentrations were higher in winter-spring (84.52 ± 40.38 µg/m³) than summer-monsoon (49.24 ± 16.96 µg/m³). UR sampler at Bhowanipore recorded 66.34 ± 6.27 µg/m³ (summer-monsoon) and 116.14 ± 53.02 µg/m³ (winter-spring). Days exceeding the NAAQS 24 h limit (60 µg/m³) were consistently greater in UR (Fig. 1b, c). X-ray Fluorescence (XRF) analysis of winter PM_2.5_ showed higher concentrations of Cr, Fe, Zn, and S in UR than RU (Fig. S1). High Performance Liquid Chromatography (HPLC) confirmed higher B(α)P in ambient PM_2.5_ samples (∼1.5× RU) and in airway cells of UR (Fig. 1d–j).

**Fig. 1:**
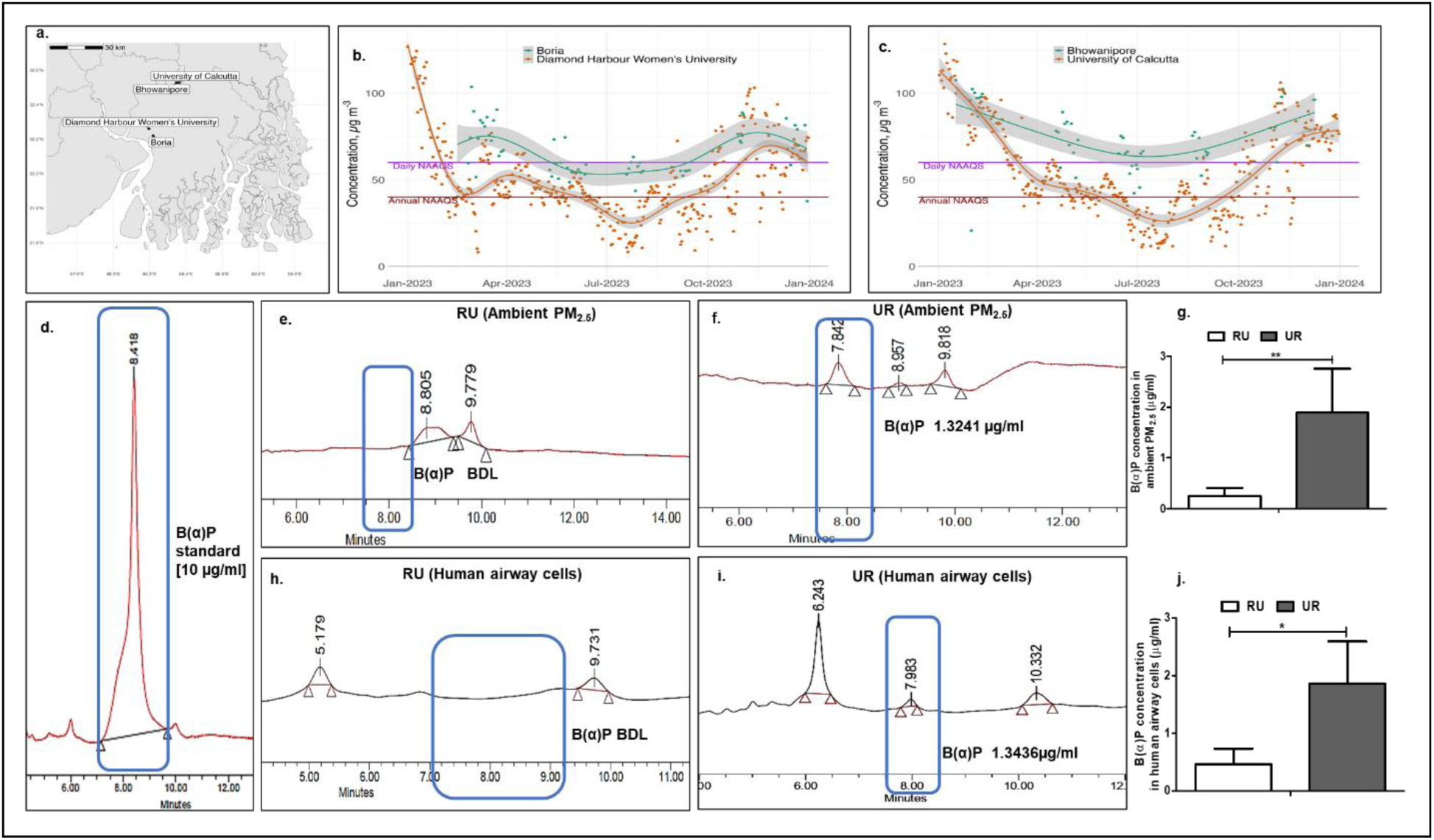
Exposure and characterization of ambient PM_2.5_ in rural and urban areas. a. Ambient PM_2.5_ concentrations observed at RU (Diamond Harbour Women’s University and Boria) and UR (University of Calcutta and Bhowanipore) locations. b. Observed daily PM_2.5_ concentrations (points) and spline fit with standard error (lines) from ground monitoring station (brown) at Diamond Harbour Women’s University and PM_2.5_ sampler (green) at Boria. c. Observed daily PM_2.5_ concentrations (points) and spline fit with standard error (lines) from ground monitoring station (brown) at University of Calcutta and PM_2.5_ sampler (green) at Bhowanipore. d. HPLC chromatogram of B(α)P standard (10µg/ml). e. B(α)P in representative samples of ambient PM_2.5_ collected from RU. f. B(α)P in representative samples of ambient PM_2.5_ collected from UR. g. Comparative mean B(α)P concentration in ambient PM_2.5_ collected from RU and UR. h. HPLC chromatogram of B(α)P in human airway cells in representative RU individual. i. HPLC chromatogram of B(α)P in human airway cells in representative UR individual. j. Comparative mean B(α)P concentration in airway cells collected from the RU and UR cohort. *[**Abbreviations:** B(α)P, Benzo(α)pyrene; BDL, Below detection limit; RU, Rural; UR, Urban;]*

### 2.2. Demography

The biological samples [RU (N, 35) and UR (N, 43)] were collected twice from the study participants-once during Jan, 2023 (winter) and second during Sep, 2023 (monsoon). The winter and monsoon samples were expected to reveal the short-term seasonal impact of ambient PM_2.5_ exposure in RU and UR cohorts. Participants were middle-aged (RU: 43.2 ± 16.15 years; UR: 33.09 ± 9.67 years), with more females in RU (71.42%) than UR (53.48%). Body Mass Index (BMI) was lower in RU (23.70 ± 4.46) than UR (25.98 ± 4.05). Socioeconomic contrasts were marked: all UR participants were graduates and either students or employed, while only 14.28% of RU were graduates and 68.57% were homemakers. Income was higher in UR (INR 85,139.00 ± 64,802.00) than RU (INR 9,971.00 ± 7,081.00). RU families were larger and predominantly permanent residents (94.28%), whereas UR participants commuted regularly (48.83% daily). Workplace proximity to roadways was similar (<50 m), but cumulative work experience was higher in RU (43.2 ± 16.15 years) than UR (5.69 ± 5.85 years). Cooking fuel use differed sharply: UR relied exclusively on Liquid Petroleum Gas (LPG) (100%), whereas RU mainly used mixed fuel-LPG and biomass (94.28%). All participants followed a non-vegetarian diet with regular fruit intake. Passive smoke exposure was more common in UR (53.48%) than RU (25.71%). Coronavirus Disease of 2019 (COVID-19) negativity (RU: 68.57%; UR: 51.16%) and mask use (RU: 80%; UR: 55.81%) were higher in RU. Upper respiratory symptoms varied, but lower respiratory symptoms showed no seasonal differences (Table S1).

### 2.3. Comparative hematology and lung function of RU and UR individuals during monsoon and winter months

Hematological parameters—including haemoglobin (Hb) (Fig. S2a), erythrocyte sedimentation rate (ESR) (Fig. S2b), total White Blood Cell (WBC) count (Fig. S2c), lymphocyte (Fig. S2d), neutrophil (Fig. S2e), eosinophil (Fig. S2f), and monocyte frequency (Fig. S2g)—remained within clinical reference ranges in both RU and UR cohorts during winter and monsoon.

In the UR cohort, significant seasonal variation was observed: Hb was reduced in winter compared to monsoon (Fig. S2a), while WBC (Fig. S2c) and lymphocyte count (Fig. S2d) were elevated. The decline in Hb may suggest increased anemia risk, consistent with previous reports linking anemia to urban PM_2.5_ exposures (Xie et al. 2022). Monocyte frequency exhibited opposite trends across cohorts, increasing in RU but decreasing in UR during winter (Fig. S2g). Intergroup analysis further revealed significantly lower monocyte levels in UR than RU in winter (Fig. S2g). Multiple linear regression analysis further demonstrated that increasing PM_2.5_ exposure was significantly associated with decreased monocyte frequency in the UR than the RU cohort during winter (β = −0.092; 95% CI: −0.172 to −0.011) (Table S2). These findings suggest a differential seasonal and spatial modulation of innate immune responses under varying exposure levels.

Pulmonary indices Forced Vital Capacity (FVC), Forced Expiratory Volume in 1 Second (FEV1) and FEV₁/FVC remained within normal limits overall. However, RU participants showed significant winter declines in FVC (Fig. S2h), FEV₁ (Fig. S2i), and FEV₁/FVC (Fig. S2j), while UR participants exhibited reduced FVC in winter (Fig. S2h). During monsoon, UR participants had significantly lower FVC (Fig. S2h) and FEV₁/FVC (Fig. S2j) compared to RU. Reduced FVC indicated restrictive physiology, whereas reduced FEV₁/FVC reflected obstructive patterns.

Overall, seasonal PM_2.5_ exposure was associated with pulmonary functional impairment in RU and hematological alterations in UR participants, reflecting differential cohort-specific responses.

### 2.4. Oxidative stress and inflammatory response in RU and UR participants during monsoon and winter months

In the RU cohort, sputum cytology showed no seasonal change in Alveolar Macrophages (AMs), but neutrophil frequency was significantly elevated in winter versus monsoon (Fig. 2a, b). In the UR cohort, both AMs and neutrophils increased during winter, and intergroup analysis revealed higher neutrophil counts in UR than RU, suggesting enhanced airway inflammation in UR, particularly in winter.

**Fig. 2:**
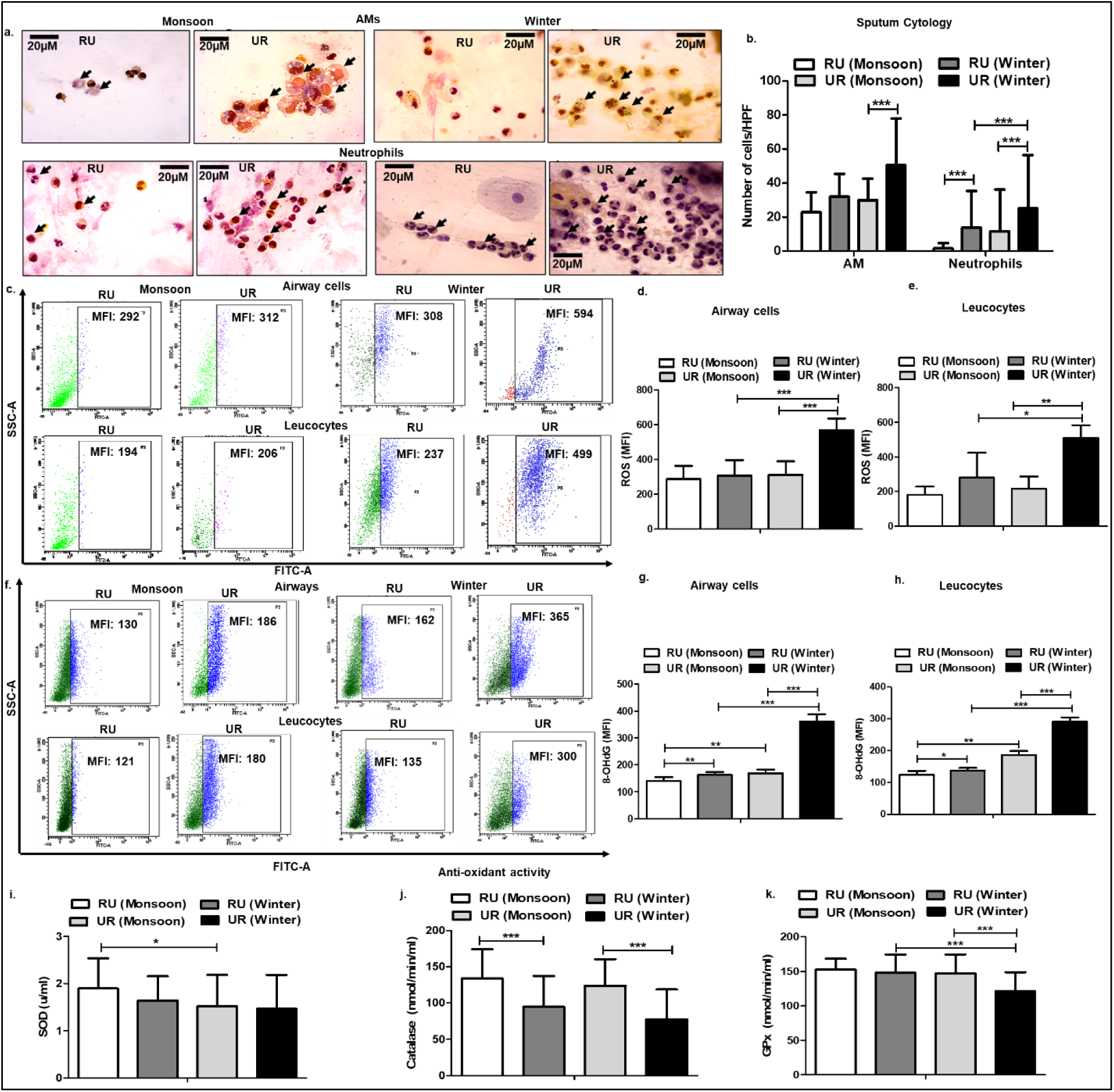
Oxidative stress and inflammatory responses in the pulmonary and the systemic microenvironments of RU and UR groups during monsoon and winter seasons [RU (N, 38); UR (N, 43); Statistical methods: LMER and MLR]. a. PAP smear of sputum showing AMs with particulate deposits and neutrophils from representative individuals of RU and UR areas (magnification 100×) during monsoon and winter b. Mean frequency of AMs and neutrophils in RU, and UR areas during two different seasons c. ROS generation in airway cells as evidenced from flowcytometry analysis of representative individuals. d. Comparative analysis of ROS production by the airway cells in the RU and the UR groups during monsoon and winter e. Comparative analysis of ROS production by the leucocytes in the RU and the UR groups during monsoon and winter f. Expression of 8-OHdG in the airway cells and the leucocytes as evidenced from flowcytometry in representative individuals g. Comparative analysis of 8-OHdG in the airway cells of the RU and the UR groups during monsoon and winter h. Comparative analysis of 8-OHdG in the leucocytes of the RU and the UR groups during monsoon and winter i. Comparative antioxidant enzyme activity of SOD from sera of RU and UR groups during monsoon and winter. j. Comparative antioxidant enzyme activity of catalase from sera of RU and UR groups during monsoon and winter. k. Comparative antioxidant enzyme activity of GPx from sera of RU and UR groups during monsoon and winter. *[**Abbreviations:** AM, Alveolar Macrophage; Gpx, Glutathione Peroxidase; LMER, Linear Mixed Effect Regression; MLR, Multiple Linear Regression; PAP, Papanicolaou; ROS, Reactive Oxygen Species; RU, Rural; SOD, Superoxide Dismutase; UR; Urban]*

Given the association between inflammation and oxidative stress, ROS levels were assessed in airway cells and leucocytes. RU participants showed no seasonal variation, whereas UR participants exhibited significantly higher ROS production in winter (Fig. 2c–e). Intergroup comparisons confirmed greater ROS levels in UR than RU during winter.

Oxidative DNA damage was evaluated by 8-Hydroxy-2’ -deoxyguanosine (8-OHdG) expression in airway cells and leucocytes (Fig. 2f–h). Both cohorts exhibited higher 8-OHdG levels in winter than monsoon. Intergroup comparisons showed consistently higher 8-OHdG in UR than RU in both cell types during both seasons (Fig. 2f–h). Moreover, multiple linear regression analysis revealed that a 1 μg/m³ increase in PM_2.5_ during the monsoon led to significantly higher 8-OHdG levels in leucocytes of UR participants compared to RU (β = 12.064; 95% CI: 3.939–20.190). Similarly, during winter, UR participants exhibited greater 8-OHdG expression in airway cells (β = 10.753; 95% CI: 2.205–13.669) and leucocytes (β = 8.346; 95% CI: 6.351–10.341) than RU participants (Table S2). Linear Mixed-Effects Regression (LMER) analysis within the UR group revealed that for every 1 μg/m³ increase in PM_2.5_ exposure, 8-OHdG levels rose significantly in airway cells (β = 5.418; 95% CI: 5.144–5.691) and leucocytes (β = 2.973; 95% CI: 2.813–3.134) (Table S3).

Antioxidant enzyme activity showed selective seasonal and cohort-specific alterations (Fig. 2i–k). Intragroup analysis revealed no significant seasonal change in Superoxide Dismutase (SOD) activity; however, intergroup comparison showed significantly reduced SOD activity in UR samples compared to RU during the monsoon (Fig. 2i). Furthermore, regression analysis indicated that each 1 μg/m³ increase in PM_2.5_ during monsoon was associated with a steeper decline in SOD levels in the UR cohort (β = −0.683; 95% CI: −1.174 to −0.192) (Table S2). Catalase activity decreased in winter across both cohorts (Fig. 2j). Glutathione Peroxidase (GPx) activity was significantly reduced in UR compared to RU during winter (Fig. 2k).

Collectively, seasonal PM_2.5_ surges, particularly in winter, contributed to increased ROS generation, oxidative DNA damage, and impaired antioxidant defence, indicating disrupted pulmonary and systemic redox balance in the UR cohort.

### 2.5. Transcriptomic analysis

As part of our earlier PM_2.5_-related health effects project, RNA sequencing [NCBI SRA: PRJNA1142236] was performed on leucocytes from asymptomatic, non-addicted, non-comorbid individuals exposed to PM_2.5_ in RU (N, 12) and UR (N, 12). Among 16,420 Differentially Expressed Genes (DEGs) identified in RU vs. UR, 319 were significantly upregulated and 92 downregulated. Kyoto Encyclopedia of Genes and Genomes (KEGG) pathway enrichment highlighted NSCLC and JAK/STAT pathways among the top 10 pathways [Table S4].

### 2.6. Bioinformatic analysis for hub genes regulating lung carcinogenesis in PM_2.5_ and B[α]P associated Lung Adenocarcinoma (LUAD) among never smokers

The Gene Expression Omnibus (GEO) dataset GSE10072, examining cigarette smoking in LUAD, comprised of never-smoker normal (N = 15) and never-smoker LUAD (N = 16) groups. Analysis identified 22,283 DEGs (padj < 0.05), including 28 upregulated and 101 downregulated genes (|Log2FC| ≥ 2). Cross-referencing with Comparative Toxicogenomics Database (CTD) revealed 9,350 DEGs shared with PM_2.5_ and B[α]P exposure (Fig. S3a). CTD Set Analyzer identified “pathways in cancer” among the top 20 enriched pathways, involving 394 genes (Table S5), which were mapped in STRING to construct a protein–protein interaction (PPI) network (Fig. S3b). Markov Cluster Algorithm (MCL) clustering defined five clusters, with the cancer pathway cluster containing 291 genes (Fig. S3c). CytoHubba analysis highlighted key regulators. The Maximal Clique Centrality (MCC) method highlighted Fibroblast Growth Factor 3 (FGF3), Epidermal Growth Factor Receptor (EGFR), Cyclin D1 (CCND1), Fibroblast Growth Factor 4 (FGF4), Catenin Beta1 (CTNNB1), proto-oncogene MYC, STAT3, Epidermal Growth Factor (EGF), Fibroblast Growth Factor 2 (FGF2), and TP53 as key hub genes (Fig. S3d). The Maximum Neighbourhood Component (MNC) algorithm identified MAPK3, Phosphatase and Tensin Homolog Deleted on Chromosome Ten (PTEN), Protein Kinase B (AKT1), EGFR, TP53, STAT3, CTNNB1, CCND1, Kirsten Rat Sarcoma Viral Oncogene Homolog (KRAS), and MYC as central nodes (Fig. S3e). Similarly, the Edge Percolated Component (EPC) method revealed STAT3, CTNNB1, TP53, EGFR, CCND1, KRAS, MYC, JUN, PTEN, and AKT1 as prominent hub genes within the cancer pathway network (Fig. S3f).

Insights from the transcriptomic and bioinformatic analyses and evidences of oxidative stress and inflammatory response in pulmonary and systemic microenvironments prompted us to investigate the JAK2/STAT3 pathway—a key stress response pathway—in the RU and UR cohorts across both monsoon and winter seasons.

### 2.7. Canonical regulation of JAK/STAT pathway in pulmonary and systemic environment during monsoon and winter seasons

The JAK/STAT pathway is closely linked to PM_2.5_-induced inflammation (Hsieh et al. 2022) To assess seasonal variation, transcript and protein levels of JAK2, STAT3, and Nuclear Factor Kappa Light Chain of Activated B Cells (NF-κB) were analyzed in airway cells (Fig. 3a, b) and leucocytes (Fig. 3c, d) of RU and UR cohorts. JAK2, STAT3 and NF-κB showed predominant upregulation in the transcript (Fig. 3a) and protein profile (Fig. 3b) of the airway cells as well as transcript (Fig. 3c) and protein profile (Fig. 3d) of the leucocytes of the UR participants especially during the winter.

**Fig. 3:**
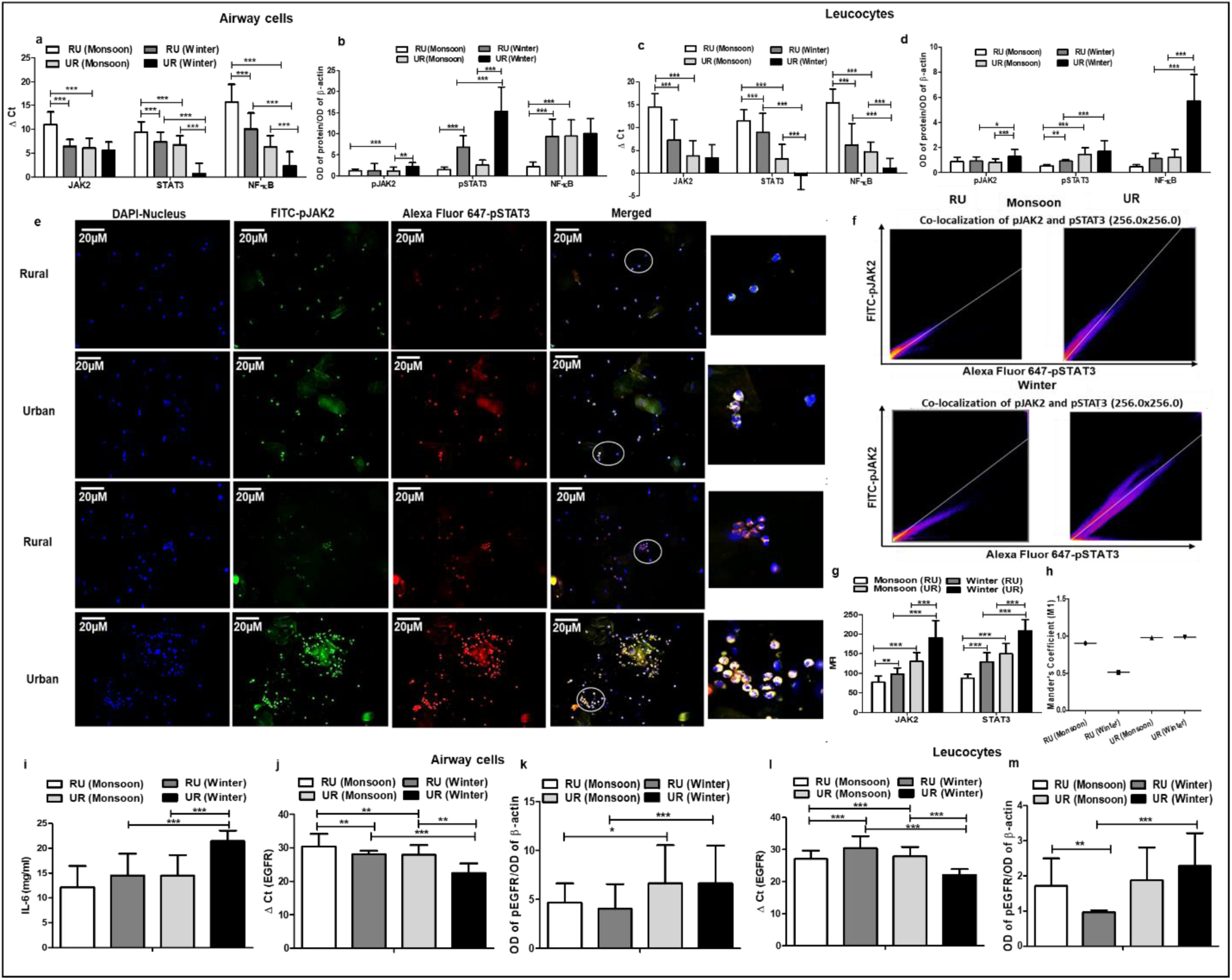
Comparative expression of oxidative stress response pathway markers (JAK2/STAT3) and its inducers in RU and UR groups during two different seasons [RU (N, 38); UR (N, 43); Statistical methods: LMER and MLR]. a. Transcript profile of JAK-2, STAT-3 and NF-κB in airway cells of RU and UR groups during monsoon and winter. b. Protein profile of JAK-2, STAT-3 and NF-κB in airway cells of RU and UR groups during monsoon and winter. c. Transcript of JAK-2, STAT-3 and NF-κB in leucocytes of RU and UR groups during monsoon and winter. d. Protein profile of JAK-2, STAT-3 and NF-κB in leucocytes of RU and UR groups during monsoon and winter. e. Co-localization of pJAK2 and pSTAT3 in airway cells of the RU and the UR cohorts during monsoon and winter as evidenced from representative images of ICC under 40× magnification f. 2D scatter plots g. MFI of JAK2 and STAT3 in RU and the UR groups during monsoon and winter h. Manders’ co-localization coefficient, M1 in RU and the UR groups during monsoon and winter i. Mean serum IL-6 of the RU and the UR cohorts during monsoon and winter j. EGFR expression in the airway cells as evidenced from transcript in RU and the UR groups during monsoon and winter. k. EGFR expression in the airway cells as evidenced from protein profile in RU and the UR groups during monsoon and winter. l. EGFR expression in the leucocytes as observed from transcript profile in RU and the UR groups during monsoon and winter. m. EGFR expression in the protein profile in RU and the UR groups during monsoon and winter. *[**Abbreviations:** EGFR, Epidermal Growth Factor Receptor; ICC, Immunocytochemistry; IL-6, Interleukin 6; JAK2, Janus Kinase 2; LMER, Linear Mixed Effect Regression; MFI, Mean Fluorescence Intensity; MLR, Multiple Linear Regression; NF-κB, Nuclear Factor Kappa Light Chain of Activated B Cells; RU, Rural; STAT3, Signal Transducer and Activator of Transcription 3; UR, Urban.]*

Multiple linear regression showed that rise in PM_2.5_ during monsoon impacted UR more than RU as observed from (i) elevated the gene expression of STAT3 (β= −1.483, 95% CI, −2.984, 0.018), NF-κB (β= −2.531, 95% CI, −4.779, −0.283) in the airway cells and (ii) JAK2 (β= −3.383, 95% CI, −5.566, −1.200), STAT3 (β= −3.350, 95% CI, −5.331, −1.369) and NF-κB (β= −3.759, 95% CI, −5.540, −1.977) in the leucocytes. Further winter PM_2.5_ surge (i) upgraded the gene expression of STAT3 (β= −0.379, 95% CI, −0.752, −0.006), NF-ᴋB (β= −0.504, 95% CI, −1.011, −0.002) within the airway cells and (ii) protein expression of NF-ᴋB (β= 0.429, 95% CI, 0.152, 0.706) within the leucocytes of the UR group more than the RU group (Table S2).

According to LMER analysis rise in PM_2.5_ during winter caused (i) upregulation of STAT3 (β= −0.170, 95% CI, −0.194, −0.145) and NF-κB gene (β= −0.112, 95% CI, −0.144, −0.081) in the airway cells and in the leucocytes [STAT3 (β= −0.099, 95% CI, −0.130, −0.067), and NF-κB (β= −0.010, 95% CI, −0.129, −0.073)]; (ii) activation of pSTAT3 in the airway cells (β= 0.358, 95% CI, 0.305, 0.412) and in the leucocytes (β= 0.007, 95% CI, −0.002, 0.016), and NF-κB expression (β= 0.127, 95% CI, 0.107, 0.146) in the leucocytes of the UR cohort (Table S3).

Phosphorylation of JAK2 (pJAK2Tyr570) and STAT3 (pSTAT3Tyr705) was greater in winter (Fig. 3e–h). UR samples showed strong pJAK2–pSTAT3 colocalization (Mander’s M1 ∼1), indicating sustained signaling, whereas RU displayed minimal overlap, suggesting STAT3 redistribution. Multiple linear regression revealed that high PM_2.5_ during winter increased both JAK2/STAT3 localization [JAK2 (β= 24.182, 95% CI, 9.655, 38.710), STAT3 (β= 17.941, 95% CI, 2.895, 32.987)] within the airway cells of UR group more than RU group (Table S2). Within the UR cohort, LMER also confirmed that pJAK2/pSTAT3 protein localization was increased in the airway cells with winter surge in PM_2.5_ [JAK2: (β= 1.720, 95% CI, 1.276, 2.164), STAT3 (β= 1.688, 95% CI, 1.335, 2.041)] (Table S3).

Interleukin-6 (IL-6), an upstream regulator, was elevated in UR serum during winter versus monsoon and higher than RU (Fig. 3i). EGFR is another upstream activator of JAK2/STAT3, particularly in Non-Small Cell Lung Cancer (NSCLC) (Zhou et al. 2022), was consistently upregulated in UR airway cells during winter (Fig. 3j–k). In leucocytes, UR showed higher winter expression of EGFR, while RU peaked in monsoon (Fig. 3l–m). To assess mutational driver, EGFR hotspot region was analyzed in sputum derived cells collected during winter, the season with peak PM_2.5_ concentrations in both regions. Mutation analysis of EGFR exon 20 (T790M), a known resistance mutation promoting enhanced ATP binding and tyrosine kinase inhibitor resistance (Bai et al. 2023) revealed no alterations (Fig. S4; GenBank accession numbers PX314130-PX314167, Table S6).

Collectively, UR participants exhibited PM_2.5_-driven IL-6 secretion and EGFR upregulation independent of EGFR exon 20 (T790M) mutation and stronger JAK2/STAT3 activation during winter.

### 2.8. Expression of the inhibitors of the JAK/STAT pathway in the RU and the UR cohorts

The JAK2/STAT3 pathway is negatively regulated by the JAK2 inhibitor, Suppressor of Cytokine Signaling 2 (SOCS2) and the STAT3 inhibitor, Protein Inhibitor of Activated STAT 2 (PIAS2) (Xin et al. 2020). We therefore assessed their molecular expression in response to seasonal PM_2.5_. The pathway was hyper-activated in UR compared to RU cohorts. Intragroup analysis showed no seasonal change in PIAS2 in RU, while SOCS2 was consistently upregulated in airway cells and leucocytes during winter (Fig. 4a, b). Intergroup analysis revealed suppression of PIAS2 in UR airway cells and leucocytes during winter, while SOCS2 was markedly reduced in UR airway cells across both seasons and in leucocytes during monsoon (Fig. 4a, b). Western blotting confirmed minimal expression of both inhibitors in UR during winter, whereas RU samples exhibited higher inhibitor expression in monsoon, except SOCS2 in leucocytes, which increased in winter (Fig. 4c–f). Multiple linear regression revealed that increase in PM_2.5_ suppressed in the gene expression of PIAS2 (β= 0.688, 95% CI, 0.097, 1.279) within leucocytes of the UR cohort more than the RU cohort during winter (Table S2). Further LMER analysis corroborated that rise in PM_2.5_ during winter within the UR cohort decreased protein expression of PIAS2 (β= −0.014, 95% CI, −0.016, −0.011) and SOCS2 (β= −0.009, 95% CI, −0.012, −0.006) in the airway cells and decreased protein expression of PIAS2 (β= −0.002, 95% CI, −0.004, −0.001) in the leucocytes (Table S3).

**Fig. 4:**
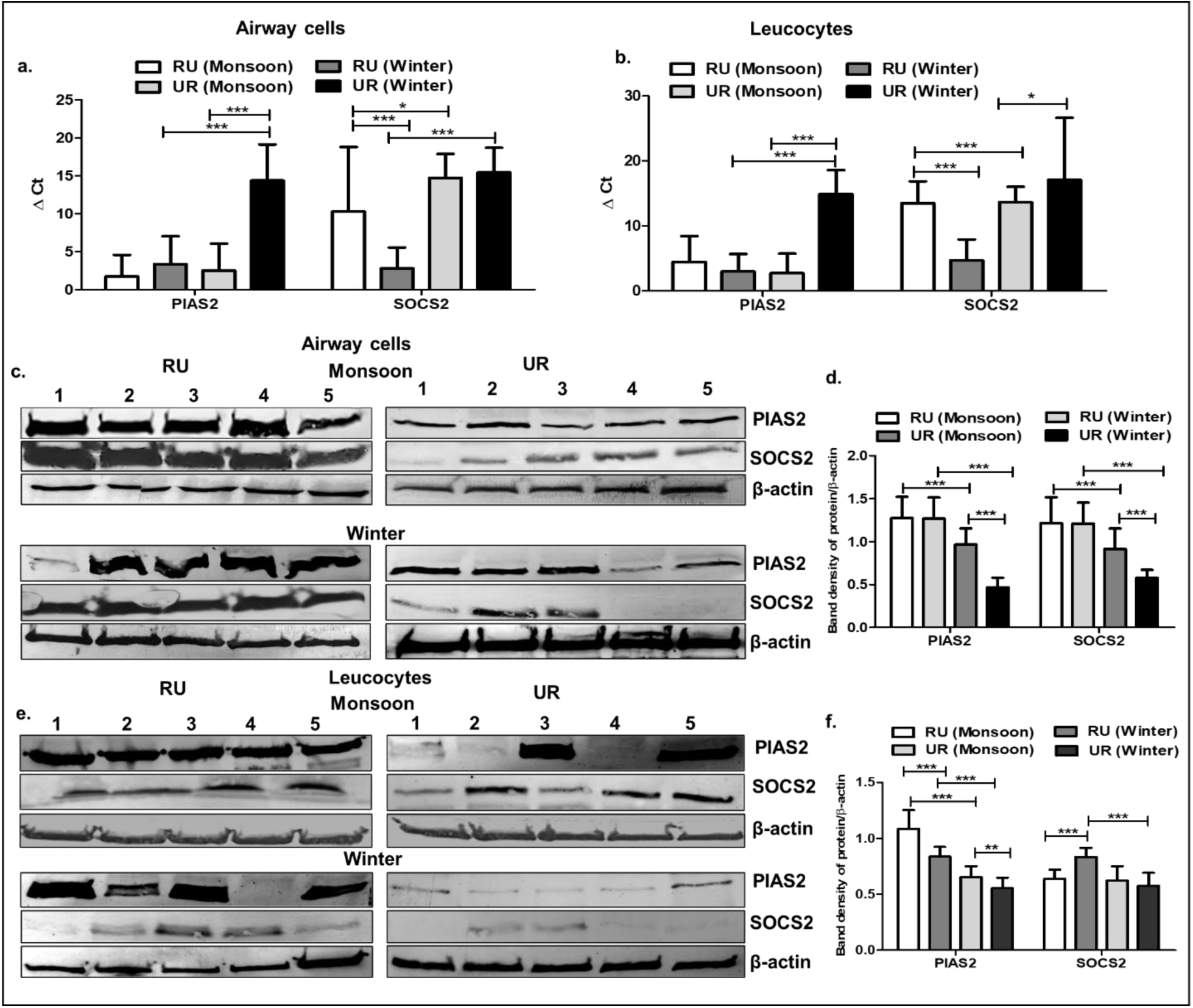
Comparative expression of JAK2 and STAT3 inhibitors in the RU and the UR groups during monsoon and winter season [RU (N, 38); UR (N, 43); Statistical methods: LMER and MLR]. a. Transcript profile of PIAS2 and SOCS2 in airway cells of RU and UR groups during monsoon and winter. b. Transcript profile of PIAS2 and SOCS2 in leucocytes of RU and UR groups during monsoon and winter. c. Representative Western blots of PIAS2 and SOCS2 during both the seasons of RU and UR groups. d. Mean band intensities in airway cells of RU and UR groups. e. Representative Western blots of PIAS2 and SOCS2 during both the seasons of RU and UR groups. f. Mean band intensities in leucocytes of RU and UR groups. *[**Abbreviations:** LMER, Linear Mixed Effect Regression; MLR, Multiple Linear Regression; PIAS2, Protein Inhibitor of Activated STAT2; SOCS2, Suppressor of Cytokine Signaling 2; RU, Rural; UR, Urban.]*

The hyper-activation of JAK2/STAT3 signaling and suppressed expression of the natural inhibitors PIAS2 and SOCS may mark a serious concern among the UR individuals who are regularly exposed to increasing amount of PM_2.5_ especially during winter.

### 2.9. Crosstalk of JAK2/STAT3 pathway with RAS-MAPK pathway in RU and UR cohorts during different seasons

Over 40% of human cancers involve hyperactivation of the Ras/Raf/ Mitogen-Activated Protein Kinase (MEK)/ Extracellular-Signal Regulated Kinase 2 (ERK) -MAPK pathway (Yuan et al. 2020). Crosstalk of JAK/STAT with Ras/Raf/MAPK has been shown in gastrointestinal cancer (Wang et al. 2021) and melanoma (Zhao et al. 2020). A PM_2.5_ dose-dependent activation of the Ras/Raf/MAPK pathway has been reported in the normal bronchial BEAS-2B cells (Wu et al. 2017). Therefore, the crosstalk of Ras/Raf/MAPK pathway with JAK2/STAT3 piqued our interest. In winter, regulatory genes Protein Tyrosine Phosphatase Non-Receptor Type 11 (PTPN11), Growth Factor Receptor-Bound Protein 2 (GRB2), KRAS and ERK2 were significantly upregulated in UR airway cells (Fig. 5a), while only PTPN11 and GRB2 increased in leucocytes (Fig. 5b). Corresponding proteins GRB2, RAS, C-RAF, p-MEK and p-ERK were robustly expressed in UR airway cells (Fig. 5c) and leucocytes (Fig. 5d).

**Fig 5:**
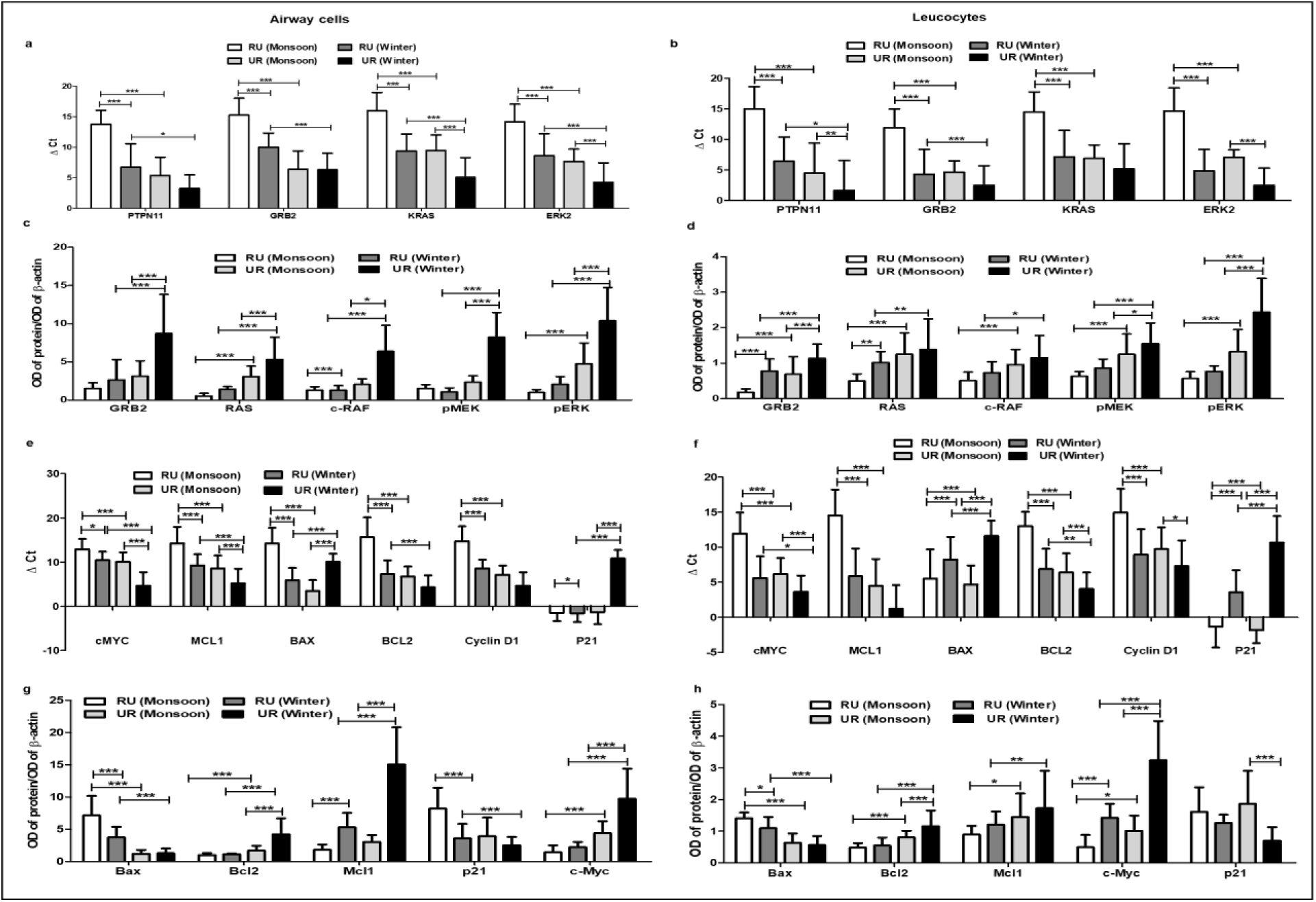
Comparative expression profile of the Ras/MAPK signaling and downstream targets in RU and the UR groups during monsoon and winter season [RU (N, 38); UR (N, 43); Statistical methods: LMER and MLR]. a. Comparative expression of PTPN11, GRB2, KRAS and ERK2 as evident from qRT-PCR in airway cells of the RU and the UR groups in two seasons. b. Comparative expression of PTPN11, GRB2, KRAS and ERK2 as evident from qRT-PCR in leucocytes of the RU and the UR groups in two seasons. c. Comparative protein expression profile of GRB2, RAS, c-RAF, p-MEK and p-ERK in the airway cells of the RU and the UR groups in the two seasons. d. Comparative protein expression profile of GRB2, RAS, c-RAF, p-MEK and p-ERK in the leucocytes of the RU and the UR groups in the two seasons. e. The comparative expression of downstream markers (c-MYC, MCL1, BAX, BCL2, cyclin D1 and p21) as evident from qRT-PCR in the airway cells of the RU and the UR groups during monsoon and winter season. f. The comparative expression of downstream markers (c-MYC, MCL1, BAX, BCL2, cyclin D1 and p21) as evident from qRT-PCR in the leucocytes of the RU and the UR groups during monsoon and winter season. g. The comparative expression of protein profile of BAX, BCL-2, C-MYC, MCL-1 and p21 in the airway cells of the RU and the UR groups during monsoon and winter season. h. The comparative expression of of protein profile of BAX, BCL-2, C-MYC, MCL-1 and p21 in the leucocytes of the RU and the UR groups during monsoon and winter season. *[**Abbreviations:** BAX, BCL2 Associated X, Apoptosis Regulator; BCL-2, B-Cell Lymphoma 2; C-RAF, RAF Proto-Oncogene Serine/Threonine-Protein Kinase; C-MYC, Cellular MYC; ERK2, Extracellular-Signal Regulated Kinase 2; GRB2, Growth Factor Receptor-Bound Protein 2; KRAS, Kirsten Rat Sarcoma Virus; LMER, Linear Mixed Effect Regression; MCL-1, Induced Myeloid Leukemia Cell Differentiation Protein; MLR, Multiple Linear Regression; p-ERK, Phosphorylated Extracellular Signal-Regulated Kinases; p-MEK, Phosphorylated-Mitogen-Activated Protein Kinase Kinase; PTPN11, Tyrosine-Protein Phosphatase Non-Receptor Type 11; QRT-PCR, Quantitative Reverse Transcription Polymerase Chain Reaction; RAS, Rat Sarcoma Virus.]*

Multiple linear regression revealed that compared to RU cohort, high PM_2.5_ in the UR cohort (i) triggered gene expression of KRAS (β= −2.336, 95% CI, −4.251, −0.421) and ERK2 (β= −2.573, 95% CI, −4.469, −0.677) in the airway cells during winter; (ii) PTPN11 (β= −4.998, 95% CI, −8.329, −1.668) and ERK2 (β= −2.948, 95% CI, −4.536, −1.361) in the leucocytes during monsoon. Additionally, increase in PM_2.5_ also upgraded the protein expression of RAS (β= 0.913, 95% CI, 0.116, 1.710) in the airway cells of the UR cohort than the RU cohort during monsoon. Similarly, during winter PM_2.5_ influenced protein expression of GRB2 (β= 0.691, 95% CI, −0.008, 1.391), pMEK (β= 0.543, 95% CI, 0.114, 0.972) in the airway cells and pERK (β= 0.151, 95% CI, 0.028, 0.274) in the leucocytes of the UR participants more than the RU participants (Table S2).

LMER analysis depicted rise in PM_2.5_ during winter (i) upregulated PTPN11 (β= −0.059, 95% CI, −0.091, −0.027), KRAS (β= −0.124, 95% CI, −0.158, −0.091), ERK2 (β=, −0.097, 95% CI, −0.130, −0.064) in the airway cells and (ii) increased gene expression of GRB2 (β= −0.061, 95% CI, −0.095, −0.027), PTPN11 (β= −0.080, 95% CI, −0.129, −0.030), KRAS (β= −0.048, 95% CI, −0.088, −0.009) and ERK2 (β= −0.130, 95% CI, −0.155, −0.105) in the leucocytes; (iii) increased protein expression of GRB2 (β= 0.158, 95% CI, 0.109, 0.207), RAS (β= 0.062, 95% CI, 0.035, 0.091), c-RAF (β= 0.122, 95% CI, 0.092, 0.152), pMEK (β= 0.166, 95% CI, 0.136, 0.196), pERK (β= 0.160, 95% CI, 0.115, 0.205) in the airway cells; (iv) GRB2 (β= 0.012, 95% CI, 0.007, 0.181) and pERK (β= 0.031, 95% CI, 0.022, 0.040) in the leucocytes of the UR cohort (Table S3).

The JAK2/STAT3 pathway regulates apoptosis and oncogenesis (Xin et al. 2020). UR airway cells (Fig. 5e) and leucocytes (Fig. 5f) showed suppressed BCL-2 Associated Protein X (BAX) and p21 but upregulated B-cell Lymphoma-2 (BCL-2), Myeloid Cell Leukemia 1 (MCL-1), c-MYC and cyclin D1. Proteins followed the same trend (Fig. 5g, h).

Multiple linear regression revealed that rise in PM_2.5_ during monsoon (i) enhanced airway gene expression of BCL-2 (β= −2.869, 95%CI, −5.255, −0.483), cyclin D1 (β= −3.265, 95% CI, −3.373, −1.576); (ii) increased leucocyte gene expression MCL-1 (β= −4.093, 95% CI, −6.868, −1.318), BCL-2 (β= −3.265, 95% CI, −3.373, −1.576) and cyclin D1 (β= −3.275, 95%CI, −5.566, −0.984) and (iii) decreased leucocyte expression of BAX (β= −0.336, 95%CI, −0.518, −0.155) in the UR more than the RU group. Similarly, during winter, PM_2.5_ rise inflicted (i) reduced protein expression of BAX (β= −0.269, 95% CI, −0.486, −0.052) in the airway cells and elevated protein expression of c-MYC (β= 0.169, 95% CI, 0.008, 0.330), BCL-2 (β= 0.065, 95% CI, −0.001, 0.133) and MCL-1 (β= 0.140, 95% CI, −0.005, 0.286) in the leucocytes of the UR subjects than the RU subjects (Table S2).

LMER analysis estimated that within the UR cohort rise in PM_2.5_ (i) increased gene expression of c-MYC (β= −0.151, 95% CI, −0.185, −0.115), BCL-2 (β= −0.069, 95% CI, −0.102, −0.037), MCL-1 (β= −0.961, 95% CI, −0.137,-0.571), cyclin D1 (β= −0.072, 95% CI, −0.103, −0.037), (ii) protein expression of c-MYC (β= 0.151, 95% CI, 0.107,0.196), BCL-2 (β= 0.072, 95% CI, 0.048, 0.096), MCL-1 (β= 0.340, 95% CI, 0.294, 0.385); (iii) decreased BAX (β= 0.186, 95% CI, 0.159, 0.214) and p21 (β= −0.039, 95% CI, −0.065, −0.012) in the airway cells; (iv) upregulated transcript profile of c-MYC (β= −0.071, 95% CI, −0.098, −0.048), BCL-2 (β=, −0.067, 95% CI, −0.093, −0.041), MCL-1 (β= −0.092, 95% CI, −0.131, −0.052), cyclin D1 (β= −0.068, 95% CI, −1.070, −0.029); and (v) protein expression of c-MYC (β= 0.063, 95% CI, 0.051, 0.074) and BCL-2 (β= 1.02, 95% CI, 5.322,1.508) in the leucocytes (Table S3).

PM_2.5_ influenced high expression of c-MYC, cyclin D1, anti-apoptotic MCL-1 and BCL2 and low expression of BAX and p21 in the UR cohort left concerns for risk of oncogenesis in the future.

### 2.10. Relative Risk (RR) for lung cancer mortality in RU and UR cohorts during monsoon and winter months

The long-term estimated risk of lung cancer mortality in the RU cohort (X₀ = 5 μg/m³) was higher in winter [RR: 1.753; 95% CI: 1.230–2.499] than monsoon [RR: 1.606; 95% CI: 1.191–2.167], reflecting a 75.3% and 60.6% increase, respectively, relative to the WHO annual PM_2.5_ guideline (5 μg/m³). In the UR cohort, risk was similarly elevated in winter [RR: 1.853; 95% CI: 1.255–2.735] compared to monsoon [RR: 1.637; 95% CI: 1.199–2.236], corresponding to increases of 85.3% and 63.7%. A similar seasonal trend was observed for short-term mortality risk (X₀ = 15 μg/m³). In RU, short-term RR was higher in winter [RR: 1.396; 95% CI: 1.131–1.723] than monsoon [RR: 1.279; 95% CI: 1.095–1.494], corresponding to 39.6% and 27.9% increases over the WHO 24 h limit (15 μg/m³). In UR, winter RR [1.475; 95% CI: 1.154–1.886] exceeded monsoon RR [1.304; 95% CI: 1.102–1.542], reflecting 47.5% and 30.4% increases, respectively.

## 3. Discussion

Indian cities are major sources of air pollution, with severe health implications for their inhabitants (Guttikunda and Ka 2022; Goel et al. 2023). Industrial activities, construction sites and vehicular emissions are primary drivers of PM_2.5_ pollution. India consistently records some of the highest ambient PM_2.5_ concentrations globally, and metropolitan areas such as Kolkata routinely exceed the annual PM_2.5_ standards set by NAAQS and WHO (Jaganathan et al. 2025). PM_2.5_ associated lung cancer, particularly among non-smokers, has emerged as a pressing concern in south east Asia including India (Myers et al. 2021; Noronha et al. 2024). In light of these facts, we aimed to investigate the effect of seasonal and spatial variations in ambient PM_2.5_ on oxidative stress response and pro-oncogenic signaling in asymptomatic individuals from urban and rural regions of West Bengal, India.

PM_2.5_ concentration and constituents diverge with different Indian seasons. Winter PM_2.5_ surges coincided with low temperatures, diminished vegetation cover (Sengupta et al. 2024), and crop residue burning (Saharan et al. 2024). Seasonal variability in PM_2.5_ toxicity has been linked to increased ROS generation and heightened pro-inflammatory cytokine release compared to summer. This is likely attributable to higher concentrations of anions (NO₃⁻, SO₄²⁻) and water-soluble metals (Al, Ca, Mg, Zn, Cr) in winter PM_2.5_ (Marchetti et al. 2019; Li et al. 2020). Consistent with these studies, we observed higher PM_2.5_ concentrations during the winter-spring season (January–March 2023) compared to the summer-monsoon period (April–September 2023) in both RU and UR areas with the increase being more pronounced in the UR area. Notably UR PM_2.5_ samples revealed higher concentrations of metals (Cr, Fe, Zn and S) associated with health risk than the RU samples. These findings resonated with previous studies that reported association of PM-bound metals (As, Cd, Co, Pb, Cr, Ni, Cu) to carcinogenic risks in Delhi (Khanna et al. 2015), Agra (Sah et al. 2017; Sah et al. 2019), Lucknow (Pandey et al. 2013), and Pune (Yadav and Satsangi 2013). Likewise, PAH such as B[α]P, a known PM_2.5_-bound carcinogen (Chang et al. 2019), was detected in UR PM_2.5_ and airway cells but were minimal in RU samples. This aligned with observations from Jharkhand coalfields and Delhi schools, where PM_2.5_-bound PAHs were associated with heightened cancer risks (Roy et al. 2019; Jyethi et al. 2014).

Hematological and spirometry evaluations revealed subtle yet significant seasonal and spatial differences between the cohorts. The UR cohort demonstrated decreased Hb levels and higher leucocyte counts in winter, indicative of systemic inflammation. Concurrently other studies also linked PM_2.5_ to anaemia in Indian children and pregnant women (Mehta et al. 2021; Xie et al. 2022). The difference in monocyte frequency between RU and UR participants during winter may be attributed to the distinct composition and immunomodulatory effects of PM_2.5_, with RU exposure triggering acute immune activation and UR exposure leading to chronic inflammation-induced monocyte depletion or tissue migration. Importantly, reductions in FVC and FEV1/FVC ratios in UR individuals during winter suggested a dual risk for restrictive and obstructive lung dysfunction. These results were coherent with global epidemiological studies linking PM_2.5_ exposure to impaired lung development (Wang et al. 2023) and lung function decrement (Mu et al. 2022). RU participants had comparatively better lung function but exhibited restrictive and obstructive tendencies during winter, likely due to biomass smoke and seasonal PM_2.5_ surges.

Oxidative stress is a well-established pathway for PM_2.5_ toxicity, implicated in DNA adduct formation, lipid peroxidation, and protein modification (Wang et al. 2024; Hou et al. 2024). Elevated ROS generation, increased 8OHdG levels, and depletion of antioxidant enzyme activities (SOD, catalase, GPx) provided direct evidence of oxidative damage in the pulmonary and the systemic compartments. The dose-dependent relationships between PM_2.5_ and oxidative stress markers in the UR cohorts reinforced the causal association. Moreover, increased neutrophilic infiltration in the UR airway samples during winter underscored a sustained inflammatory state, which may act synergistically with oxidative stress to drive lung tissue injury (Li et al. 2023).

The impact of PM_2.5_ on the oxidative stress markers and proinflammatory response was further reinstated by the transcriptomic analysis that revealed perturbations in the stress response pathway-JAK/STAT signaling (Liu et al. 2018; Liu et al. 2022). Bioinformatic validation identified hub genes (MAPK, AKT, PTEN, EGFR, STAT3, MYC, KRAS) associated with PM_2.5_-and B[α]P-linked LUAD in never-smokers. The JAK/STAT pathway is a central mediator of inflammatory responses and has been implicated in PM_2.5_-induced inflammatory response, oxidative stress and lung fibrosis in pre-clinical models (Xu et al. 2020; Yue et al. 2021). IL-6 mediated activation of JAK2/STAT3 pathway has been well evidenced in a spectrum of cancers including liver, breast, colorectal, gastric, lung, pancreatic and ovarian cancer (Huang et al. 2022). However, evidence linking JAK/STAT signaling to oxidative stress and pro-carcinogenic alterations in high PM_2.5_ exposed asymptomatic Indian populations remains limited. Our findings demonstrated that the winter elevation in PM_2.5_ concentrations activated IL-6/JAK2/STAT3 signaling pathways in the pulmonary and systemic compartments of both RU and UR cohorts, with a markedly greater impact on the UR cohort. EGFR activation has been well established with Programmed Death-Ligand 1 mediated therapeutic resistance through upregulation of IL-6/JAK2//STAT3 mediated in NSCLC (Zhang et al. 2016). Exposure to PM_2.5_ was linked to the presence of EGFR (18%) and KRAS (53%) driver mutations in histologically normal lung tissue samples (Hill et al. 2023). Lung cancer is often characterized by EGFR overexpression and mutations in exon 20, both of which critically influence therapeutic outcomes (O’Sullivan et al. 2022). Though we observed EGFR overexpression in UR during winter surge in PM_2.5_, we did not observe any significant changes in mutational profile of exon 20 of EGFR.

PM_2.5_ exposure has been shown to activate the Ras/Raf/MAPK pathway in a dose-dependent manner in normal human bronchial epithelial BEAS-2B cells (Wu et al. 2017), highlighting its potential role in initiating pro-carcinogenic signaling in airway epithelium. Importantly, therapeutic crosstalk between the JAK/STAT and Ras/Raf//MAPK pathways has been demonstrated in gastrointestinal cancers and melanoma, where simultaneous inhibition of JAK2/STAT3 and ERK signaling suppressed tumor proliferation and induces apoptosis (Wang et al. 2021; Zhao et al. 2020). Furthermore, PTPN activates KRAS and downstream signaling which is crucial for survival and proliferation of KRAS driven cancer cells (Huang et al. 2020). GRB2 is an adaptor protein that forms stable complexes-mediated via SH2 domain-with tyrosine phosphorylated EGFR (Lowenstein et al. 1992) leading to the activation of RAS and its downstream kinases, ERK1/2. In congruence with these reports, we observed concomitant upregulation of Ras/Raf/MAPK pathway (PTPN11, GRB2, KRAS, ERK2) and their downstream effectors (c-MYC, BCL-2, MCL-1, cyclin D1) in the airway cells and leucocytes of the RU and UR cohorts especially with decline of air quality during winter. This suggested that PM_2.5_ exposure not only triggers an acute stress response but also promotes a microenvironment conducive to malignant transformation. Notably, suppression of tumor suppressors (p21, BAX) and JAK2/STAT3 inhibitory regulators -SOCS2, PIAS2 (Xin et al. 2020) in UR individuals likely contributed to unchecked signaling activity. This aligned with studies that demonstrated that the negative regulators of JAK/STAT pathway, SOCS2 (Ma et al. 2023) and PIAS genes (Zhang et al. 2024) were downregulated in NSCLC.

The elevated RR of lung cancer mortality in UR cohorts, particularly during winter, underscored the public health relevance of these findings. Long-term RR estimates suggested higher risk of lung cancer mortality among UR individuals compared to WHO guideline levels, echoing global reports linking ambient PM_2.5_ to lung cancer incidence in never-smokers (Myers et al. 2021; Noronha et al. 2024). The observed short-term RR elevation further highlighted the acute health burden during episodic PM_2.5_ surges.

This study comes along with a few limitations. First, its observational design limited the ability to establish direct causal relationships. Second, the relatively small sample size for molecular analyses might have affected generalizability. Third, reliance on self-reported data and ambient PM_2.5_ levels as proxies for personal exposure, the potential influence of unmeasured PM_2.5_ constituents, and residual confounding factors further constrained interpretation. Future studies may adopt longitudinal designs with larger cohorts and high-resolution exposure assessments to strengthen causal inference. Nonetheless, the consistent associations observed between real-time PM_2.5_ levels and biomarkers enhanced robustness of our findings. Importantly, the study’s strength lied in its integrative approach, combining environmental, cellular, and molecular data to comprehensively delineate PM_2.5_-related oxidative stress and pro-oncogenic signaling.

It may be summarized that seasonal and spatial differences in ambient PM_2.5_ exposure may drive distinct pro-carcinogenic changes in asymptomatic urban and rural cohorts, with a greater impact in the urban cohort. Elevated PM_2.5_ concentrations during winter triggered oxidative stress, evidenced by increased ROS and 8-OHdG levels and depletion of antioxidants (SOD, catalase, GPx), alongside airway inflammation. At the molecular level, activation of the IL-6 or EGFR mediated JAK2/STAT3 axis and crosstalk with Ras/Raf/MAPK signaling promoted pro-survival and proliferative phenotypes, marked by upregulation of BCL-2, MCL-1, c-MYC, and cyclin D1, and suppression of tumor suppressors (BAX, p21). Notably, reduced SOCS2 and PIAS2 expression likely sustained JAK2/STAT3 hyperactivation. Together, these findings highlighted the interplay between PM_2.5_-induced oxidative stress and JAK2/STAT3 mediated pro-oncogenic signaling in pulmonary and systemic compartments, emphasizing the need for air quality interventions, molecular surveillance and targeted mitigation strategies in high-exposure populations.

## 4. Materials and methods

### 4.1. Study areas and ambient PM_2.5_ exposure analysis

The study areas included an urban area of Bhowanipore, Kolkata (UR) and a rural area of Boria, Diamond Harbor (RU) in West Bengal, India. The extent of ambient PM_2.5_ exposure at UR (coordinates: 22.52501°N, 88.34667°E) were estimated by a PM_2.5_ sampler [APM 550EL, Envirotech, New Delhi, India]. Additionally, simultaneous ambient PM_2.5_ data was also obtained from the nearest (2 km from UR site) air quality monitoring station of West Bengal Pollution Control Board (WBPCB), placed at Taraknath Palit Siksha Prangan (Coordinates: 22.3137 °N 88.2146 °E) (https://www.wbpcb.gov.in). Similarly, at the RU (Coordinates: 22.22039 °N, 88.21917 °E), ambient PM_2.5_ data was collected by a PM_2.5_ sampler and additional ambient PM_2.5_ data was obtained from WBPCB’s ambient air quality monitoring station placed at Diamond Harbor Women’s University (Coordinates: 22.26088 °N, 88.19616 °E), 12.5 km away from RU site. Since the study aimed to assess the impacts of seasonal variation of ambient PM_2.5_ on human health, PM_2.5_ data collection was conducted from Jan, 2023 to Dec, 2023.

### 4.2. Element analysis by XRF

PM_2.5_ was collected on polytetrafluoroethylene (PTFE) filter papers at RU and UR sites using PM_2.5_ samplers. Elemental concentrations in PM_2.5_ were determined using PTFE membrane filters (Cat#7592-104) (Whatman, Cytiva, Marlborough, MA, USA) analyzed by XRF on a HORIBA Scientific XGT-7200 X-ray Analytical Microscope (Kyoto, Japan). Measurements were performed at the University Grants Commission (UGC)-Department of Atomic Energy (DAE) Consortium for Scientific Research, Kolkata. XRF analysis was performed with detectors set at 50 kV, 1 mA, XGT 1.2 mm, and a titanium (Ti) filter, with a preset acquisition time of 1800 s. Calibration of the XRF instrument was performed using a copper (Cu) standard.

### 4.3. Study participants

Subjects from the study areas were enrolled after obtaining informed consent. Inclusion criteria comprised apparently healthy, non-addicted (asymptomatic) adult males and females aged 25–60 years, with a minimum of 8–10 h of daily occupational exposure in the study areas at least for the past five years. Exclusion criteria included individuals with comorbidities, those under medication, users of oral contraceptives, individuals with a history of malignancy, pregnant or lactating women, smokers/ex-smokers, and persons with other forms of addiction (chewing tobacco, betel nut/pan, alcohol, etc.). Screening was performed using structured questionnaires. At the UR site, 157 individuals were screened, with 43 consenting to participate in both seasonal assessments. At the RU site, 90 individuals were screened, of whom 35 consented to participate in both seasons. The study was conducted in accordance with the Declaration of Helsinki and approved by the Institutional Ethics Committee (CNCI-IEC-DS-2019-10, dated 23 July 2019).

### 4.4. Sample collection

Blood samples (3 ml) were collected from ante-cubital vein in vacutainer tubes (BD Biosciences, Franklin Lakes, NJ, USA) with and without K_2_EDTA as anticoagulant for whole blood and serum samples respectively. The spontaneously expectorated sputum in early morning hours was collected from each participant and was separated into two parts-(i) in a tube with dithiothreitol (DTT) (Cat#D0632) (Merck, Rahway, NJ, USA) as a mucolytic agent and (ii) air-dried smears on slides.

### 4.5. Hematology and lung function test

Hematological analysis was performed using an automated hematology analyzer (Erma PCE-210, Saitama, Japan). Pulmonary function tests (PFTs) were conducted with PC spirometry (SP260, Schiller, Barr, Switzerland) following a published protocol (Das et al. 2014). Measured values were compared with predicted values adjusted for age, gender, height, and ethnicity. Parameters recorded for analysis included FVC, FEV1 and the FEV1/FVC ratio.

### 4.6. Sputum cytology

Sputum cytology was assessed using Papanicolaou (PAP) staining. Cell counts (excluding squamous epithelial cells) were performed across 10 randomly selected fields under a light microscope (CX40, Olympus, Tokyo, Japan). Samples with <20% squamous cell contamination and presence of cylindrical epithelial cells and/or AMs were deemed adequate. AMs, bronchial epithelial cells, neutrophils, eosinophils, lymphocytes, and goblet cells were identified following established criteria (Herbert 1989).

### 4.7. Isolation of airway cells, leucocytes and protein estimation

Whole blood was mixed with solution A [0.87% NH₄Cl in 10 mM Tris–HCl, pH 7.2] at a 1:3 ratio, incubated on ice for 30 min, and centrifuged at 400×g for 20 min. The supernatant was discarded, and the process was repeated for the pellets. The resulting pellet was resuspended in solution B [0.25 mM meso-inositol, 1 mM MgCl₂ in 10 mM Na₂PO₄ buffer, pH 7.2] and centrifuged at 1200 rpm for 6 min. The final pellet consisted of leucocytes.

Sputum samples were treated with DTT and centrifuged at 1000×g for 5–6 min at room temperature. The clear supernatant was discarded and the remaining viscous fraction was further centrifuged (7000×g, 5–6 min) with Phosphate-Buffered Saline (PBS) to pellet airway cells, which were filtered through 40 µm cell strainers. Airway cells (neutrophils, eosinophils, leukocytes, AMs, and epithelial cells) were cryopreserved. For downstream analyses, RNA, DNA, and protein were extracted from these cryopreserved cells using a bead-bashing protocol with ZR BashingBead Lysis Tubes (Cat#S6003-50) (0.1 mm and 0.5 mm; Zymo Research, Irvine, CA, USA). This was done to exclude bacterial contamination according to the manufacturer’s instructions

Leucocyte and airway cells were lysed with Radioimmunoprecipitation Assay Buffer (RIPA) according to previous published protocol (Prasad et al. 2023) and cell extract was further subjected to protein estimation according to Lowry’s method (Lowry et al. 1951).

### 4.8. Estimation of PAHs by HPLC

Filters with ambient PM_2.5_ were desiccated for 24h, extracted with HPLC grade water and sonicated for 1h. The extract was then filtered and lyophilized using LaboGene Scanvac CoolSafe 4L Freeze Dryer (Allerød, Hovedstaden, Denmark). The lyophilized PM_2.5_ was dissolved in PBS for HPLC analysis.

Airway cells obtained from sputum were washed with PBS, pelleted, and lysed in acetonitrile by sonication on ice. The lysates were centrifuged at 14,000 × g, and the resulting supernatants were filtered and directly used for HPLC analysis.

HPLC analysis was performed using Waters Nova-Pak reversed-phase C18 column (3.9 mm × 150 mm, 4μm; Part No: 086344) to determine the PAHs like benzo(α)pyrene (BαP) present in the filter papers as well as in the airway cells. The samples were analysed using a gradient elution of water (A) and acetonitrile (B) with a ratio from 80(A):20(B) to 20(A):80(B). A constant flow rate of 0.1 mL/min was maintained at 254 nm. The samples were maintained at room temperature (RT) and the injection volume was 20μL.

### 4.9. Study of ROS generation

The ROS levels were determined in leucocytes and airway cells by 2’,7’-dichlorodihydrofluorescein diacetate (Cat#D399) (DCFH-DA) (Thermo Fisher Scientific /Invitrogen, Carlsbad, CA, USA) in accordance with the manufacturer’s protocol. In brief, the cell suspension was mixed with diluted DCFH-DA reagent (10 µM) and incubated at 37°C for 45 min. ROS generation was detected by flow cytometer-LSR Fortessa (BD, Franklin Lakes, NJ, USA) and analyzed with FACSDiva™ (BD, Franklin Lakes, NJ, USA).

### 4.10. Study of 8-OHdG

Leucocytes and airway cells were washed with PBS and fixed using paraformaldehyde solution (2%) for 10 min at 37 °C. The cells were then washed and permeabilized with Triton-X-100 (0.5%) for 15 min at RT. Next the cells were blocked with bovine serum albumin (BSA) (5%) for 1 h. Subsequently, the cells were washed and incubated with anti-8-OHdG antibody (Cat#sc-66036) (Santa Cruz, Dallas, TX, USA) overnight at 4°C. Finally, acquisition of the samples was done in LSR Fortessa and analyzed with FACSDiva™ (BD, Franklin Lakes, NJ, USA)

### 4.11. Study of anti-oxidant activity

The activities of GPx (Cat#703102), catalase (Cat#707002), and SOD (Cat#706002) were quantified using commercial kits (Cayman Chemical, Ann Arbor, MI, USA). GPx activity was measured in cell lysates, while catalase and SOD activities were assessed in serum.

#### 4.11.1. Sample preparation

Cell lysates were obtained by centrifugation at 4 °C for 5 min, followed by washing and centrifugation at 10,000 g for 15 min at 4 °C. Supernatants were collected for GPx assay. Serum was separated from non-EDTA blood tubes by centrifugation for SOD and catalase assays.

#### 4.11.2. SOD assay

Wells of a 96-well plate were loaded with diluted radical detector and either standard or sample, followed by xanthine oxidase to initiate the reaction. Plates were covered, incubated with shaking for 30 min at RT and absorbance was read using a microplate reader at 450 nm (Infinite® 200 PRO, TECAN, Männedorf, Zürich, Switzerland).

#### 4.11.3. GPx assay

Background wells contained assay buffer, co-substrate mixture, and NADPH; positive controls additionally contained diluted GPx control. Sample wells contained assay buffer, co-substrate mixture, NADPH, and sample. Reactions were initiated with cumene hydroperoxide, briefly shaken, and absorbance was measured at 340 nm every minute for 5 min using the microplate reader as above.

#### 4.11.4. Catalase assay

Assay buffer and methanol were added to wells, followed by catalase standards or samples. Reactions were initiated with H₂O₂ and incubated for 20 min at RT with shaking. Purpald reagent (4-amino-3-hydrazino-5-mercapto-1,2,4-triazole) and H₂O₂ were then added and incubated for 10 min, followed by potassium periodate. After 5 min shaking, absorbance was recorded at 540 nm using the microplate reader as above.

### 4.12. Assessment of pro-inflammatory cytokine

Enzyme-Linked Immunosorbent Assay (ELISA) was performed for IL-6 according to the manufacturer’s instructions (Cat#ELH-IL6-1) (RayBiotech, Norcross GA, USA). Serum was added to 96-well plates after dilution and incubated overnight at 4°C. The plates were thoroughly washed and incubated with biotin-conjugated antibody for 2 h. Subsequently wells were again washed, incubated with Horseradish peroxidase (HRP)-conjugated streptavidin, and finally added with 3,3’,5,5’-tetramethybenzidine (TMB) One-Step Substrate Reagent. The Optical Density (OD) value was measured using a microplate reader at 450 nm as already stated.

### 4.13. RNA sequencing and data analysis

Blood was collected in PAXgene Blood RNA Tubes (PreAnalytiX; Hombrechtikon, Switzerland) from RU (N,12) and UR (N,12) individuals and processed at the National Institute of Biomedical Genomics (NIBMG), India. Total RNA was extracted from leukocytes using TRIzol™ Reagent (Cat#15596026) (Thermo Fisher Scientific/Ambion; Carlsbad, CA, USA). Ribosomal RNA was depleted, and libraries were prepared with the TruSeq Stranded Total RNA kit (Illumina, San Diego, CA, USA). Paired-end (2×100 bp) sequencing was performed on a NovaSeq™ 6000 platform (Illumina, Inc., San Diego, CA, USA). Raw FASTQ files were aligned to the NCBI reference genome using the DRAGEN RNA aligner (Illumina, San Diego, CA, USA). Normalization and differential gene expression analysis were performed with DESeq2. Genes with |log2FC| ≥ 1.5 and adjusted p-value (padj) < 0.05 were considered significant. Enrichment analyses for Gene Ontology (GO) and KEGG pathways were performed using ShinyGO v0.76, and the top twenty enriched KEGG pathways were reported.

### 4.14. Bioinformatic identification of hub genes driving PM_2.5_-associated LUAD among never-smokers

GEO datasets were retrieved from the NCBI GEO repository (https://www.ncbi.nlm.nih.gov/). Differential gene expression analysis was performed using GEO2R, and the identified differentially expressed genes were compared with PM_2.5_- and B[α]P-associated gene sets from CTD (https://ctdbase.org/) using MyGeneVenn. The overlapping genes were further analyzed in CTD for pathway enrichment. Selected pathways were queried in STRING v12.0 (https://string-db.org/) to construct PPI networks. The PPI networks were clustered using the MCL clustering to identify natural groupings based on stochastic flow. Key clusters were further analyzed in Cytoscape v3.10.3 using the CytoHubba plugin to identify hub genes.

### 4.15. Real-time Quantitative Polymerase Chain Reaction (RT-qPCR)

Total RNA was extracted from airway cells and leukocytes using the PureLink™ RNA Mini Kit (Cat#12183018A) (Thermo Fisher Scientific, Carlsbad, CA, USA). RNA quality and quantity were assessed and cDNA was synthesized using the RevertAid RT Reverse Transcription Kit (Cat#K1691) (Thermo Fisher Scientific, Carlsbad, CA, USA) with 2 µg of RNA. Differential gene expression was analyzed using FastStart Essential DNA Green Master (Roche Diagnostics, Rotkreuz, Switzerland) and gene-specific primers [JAK2, STAT-3, NF-κB, EGFR, PTPN11, KRAS, GRB2, ERK2, c-MYC, MCL-1, p21, BCL-2, BAX, cyclin D1, PIAS2, SOCS2 and glyceraldehyde 3-phosphate dehydrogenase (GAPDH), Table S7] on a LightCycler® 96 (Roche Diagnostics, Rotkreuz, Switzerland). PCR conditions included pre-incubation at 98 °C for 1 min; 45 cycles of denaturation (98 °C, 10 s), annealing (50–60 °C, 40 s), and elongation (72 °C, 30 s); followed by a final extension at 72 °C for 5 min. Relative gene expression was determined using the comparative threshold cycle (ΔΔCt) method. All reactions were performed in triplicate.

### 4.16. Protein expression detection by ELISA

Proteins isolated from airway cells and leukocytes (10 µg/well) were coated onto 96-well plates with PBS and incubated overnight at 4 °C in a humid chamber. Wells were washed with phosphate buffer saline-Tween 20 (PBST) and blocked with 1% BSA for 1 h at room temperature with shaking to prevent nonspecific binding. After washing, samples were incubated with primary antibodies-JAK2, pJAK2Tyr1007 (Cat# ITP0155-50u) (G-Biosciences, St Louis, MO, USA), pSTAT3Tyr705 (Cat#sc-81523), BAX (Cat#sc-70408), BCL-2 (Cat#sc-130307), p21 (Cat#sc-397), pMEK1/2 (ser218/ser222) (Cat# sc-7995), pERK1/2 (thr202/tyr204) (Cat# sc-7383), c-MYC (Cat#4233) (Santa Cruz Biotechnologies, Dallas, TX, USA), NF-κB (Cat#BF8005), GRB2 (Cat# DF4092), RAS (Cat#5175992F2), c-RAF (Cat# AF6065), MCL1 (Cat#AF5311) (Affinity Biosciences, Cincinnati, OH, USA) and pEGFRtyr1068 (Cat# 2234) (Cell Signalling Technology Danvers, MA, USA), for 2 h at RT and subsequently incubated with HRP-conjugated secondary antibodies for 1 h at RT. Wells were washed, incubated with TMB (Cat#T4444) (Merck, Rahway, NJ, USA) substrate for 30-45 min until color development, and the reaction was stopped 1M H_2_SO_4_. OD was measured at 450 nm using a microplate reader as before.

### 4.17. Immunocytochemistry (ICC)

JAK2 and STAT3 localization in airway cells was analyzed by ICC. Sputum smears were fixed in chilled methanol (20 min, −20 °C), washed with PBS with Tween 20 (PBST) (×3), and treated with acetone for 10 min with shaking. Subsequently, slides were blocked with 5% BSA for 1.5 h at RT. Primary antibodies (JAK2 (Cat#sc-390539), STAT3 (Cat#sc-8019) (Santa Cruz Biotechnologies, Dallas, TX, USA), 1:1000 dilution in 1% BSA/PBS) were applied overnight at 4 °C in a humid chamber. Following PBST washes, samples were incubated with Fluorescein Isothiocyanate (FITC) (Cat# sc-65218) (Santa Cruz Biotechnologies, Dallas, TX, USA)- and Alexa Fluor 647 (Cat#ab150075) (Abcam, Cambridge, UK)-conjugated secondary antibodies (1:2000 dilution in 1% BSA/PBS) for 2 h at RT. Nuclei were counterstained with 4′,6-diamidino-2-phenylindole (DAPI) (Cat#D9542) (Merck, Rahway, NJ, USA), washed, mounted with glycerol, and visualized under a fluorescence microscope (MVX10, Olympus, Tokyo, Japan). Quantification was done by Image J Fiji Image Processing Software [National Institute of Health (NIH), September 14, 2022].

### 4.18. Western blot

Proteins isolated from airway cells and leucocytes were resolved on 12% Sodium Dodecyl Sulphate-Polyacrylamide Gel Electrophoresis (SDS-PAGE) and transferred onto Polyvinylidene Fluoride (PVDF) membranes. Following transfer, membranes were blocked with 5% BSA and incubated overnight at 4 °C with primary antibodies against PIAS2 (Cat#AF0273) and SOCS2 (Cat#DF8133) (Affinity Biosciences, Cincinnati, OH, USA) and β-actin (Cat#AC026) (Abclonal, Woburn, MA, USA). After washing, membranes were incubated with alkaline phosphatase (AP)-conjugated secondary antibodies for 2 h at RT. Protein bands were visualized using 5-Bromo-4-Chloro-3-Indolyl Phosphate/Nitro-blue Tetrazolium Chloride (BCIP®/NBT) (Cat#203790) substrate solution (Merck, Rahway, NJ, USA), and band intensities were quantified using ImageJ software (NIH, Bethesda, MD, USA).

### 4.19. Study of Exon Mutations of EGFR

#### 4.19.1. Preparation of Genomic DNA

The sputum derived cell pellet obtained from bashing bead method was suspended in cell lysis buffer containing proteinase K, RNase A, and 10% SDS, followed by overnight incubation at 37 °C. The lysate was then mixed with an equal volume of Tris-saturated phenol, gently agitated for 10–15 min, and centrifuged. The aqueous phase was collected and subjected to further extraction with Tris-saturated phenol, Chloroform:Isoamyl alcohol, and RNase H treatment, followed by centrifugation. The final aqueous phase was precipitated with chilled sodium acetate and isopropanol, incubated overnight at −20 °C, centrifuged, and the pellet washed with chilled ethanol. The dried pellet was resuspended in Tris-EDTA buffer for downstream applications. DNA concentration and purity were measured using a NanoDrop™ 2000 spectrophotometer (Thermo Fisher Scientific, Waltham, MA, USA).

#### 4.19.2. EGFR Mutation Analysis by Direct Sanger Sequencing

EGFR exon 20 (T790M point mutation) were analyzed by direct Sanger sequencing. Genomic DNA (100 ng) was amplified and purified using ExoSAP-IT™ (Cat#15513687) (Affymetrix, Cleveland, OH, USA) and further subjected to cycle sequencing with the BigDye® Terminator v3.1 Cycle Sequencing Kit (Cat#4337455) (Thermo Fisher Scientific, Waltham, MA, USA). Post-sequencing cleanup was performed using the NucleoSEQ® Kit (Cat#12778462) (Macherey-Nagel, Düren, Germany). Samples were eluted in 20 μL Hi-Di™ Formamide (Cat#4311320) (Thermo Fisher Scientific, Waltham, MA, USA), denatured at 95 °C for 1 min, and analyzed on an Applied Biosystems® 3130XL Genetic Analyzer (Thermo Fisher Scientific, Waltham, MA, USA). Sequence chromatograms were processed using the BioEdit Sequence Alignment Editor (Ibis Biosciences, Carlsbad, CA, USA). The details of the primer used had been enclosed in Table S8.

### 4.20. Statistical Analysis

Descriptive statistics were used to summarize the data. Associations between hematological and respiratory parameters and PM_2.5_ exposure were assessed using Pearson’s chi-square test, with p < 0.05 considered statistically significant. Transcript and protein expression levels across monsoon and spring were compared using paired t-tests, with Bonferroni correction applied for multiple comparisons (α=0.05). Multiple linear regression was also performed to evaluate associations between dependent variables (mRNA/protein expression, hematology, PFT, sputum cytology, 8-OHdG, cytokines) and independent variables, including PM_2.5_ exposure and socio-demographic covariates across RU and UR cohorts during winter and monsoon. LMER models were employed within each cohort (RU/UR) to assess the seasonal effect of PM_2.5_ exposure on hematological parameters, pulmonary function, sputum cytology, and differentially expressed genes/proteins. Covariates included age, BMI, COVID-19 infection status, marital status, household size, education, profession, income, work duration, number of working years, transportation mode, passive smoking exposure, and type of cooking fuel. Participant identity was included as a random effect in the LMER model. The RR of lung cancer mortality due to long term exposure of PM_2.5_ was calculated according to the formula RR = [(X + 1)/ (X_0_ + 1)]^β^ (Ostro 2004) where X represented mean PM_2.5_ during monsoon/spring, long term X_0_ was the WHO annual mean (5 µg/m³) and short term X_0_ was WHO 24-h average (15 µg/m³) PM_2.5_ standards (WHO, 2024), and β = 0.23218 (Ostro 2004). Statistical analyses were conducted using R v4.3.3, SPSS v25.0 (IBM, NY, USA), and GraphPad Prism v5.0 (Dotmatics, Boston, MA, USA).

## Data Availability

The datasets produced in this study are available in the following databases:

- RNA-Seq data: National Centre for Biotechnology Information (NCBI) Sequence Research Archive (SRA) (PRJNA1142236) https://www.ncbi.nlm.nih.gov/sra/?term=PRJNA1142236
- Sanger seq data of EGFR exon 20: GenBank accession numbers PX314130-PX314167, Table S6

## Acknowledgement

The authors are indebted to West Bengal Pollution Control board (Grant No. 3907–1M– 24/2010(Part–III) providing fellowships to SG and RC, as well as the project grant to DS. The authors are grateful to the National Institute of Biomedical Genomics (NIBMG), Kalyani for conducting RNA sequencing, to Dr. Sandeep Sitaram Ghugre, University Grants Commission-Department of Atomic Energy Consortium for Scientific Research (UGC-DAE CSR), Kolkata for his expertise while performing XRF and to Bose Institute, Kolkata for performing DNA Sanger Sequencing. The authors are thankful to the subjects who volunteered to participate in this study and to Director, Chittaranjan National Cancer Institute, Kolkata for the infrastructural support.

## The paper explained Problem

Although PM_2.5_ exposure is linked to lung cancer risk in both smokers and non-smokers, the early molecular alterations in asymptomatic individuals—particularly in the Indian context of seasonal fluctuations and rural-urban disparities in exposure, pollutant composition, and susceptibility—remain poorly understood. This study investigated the seasonal and spatial variations of PM_2.5_ exposure in rural and urban cohorts of West Bengal, India with a focus on oxidative stress and oncogenic signaling.

## Results

Winter monitoring showed higher PM_2.5_ and associated B[α]P in urban areas is driving oxidative stress, antioxidant depletion, and airway inflammation. Transcriptomic and bioinformatic analyses revealed IL-6/EGFR-mediated JAK2/STAT3 activation and Ras/Raf/MAPK crosstalk, marked by upregulation of pro-survival genes (BCL-2, MCL-1, c-MYC, cyclin D1), suppression of tumor suppressors (BAX, p21), and loss of JAK/STAT negative regulators (PIAS2, SOCS2), sustaining oncogenic signaling. Statistical modeling further linked winter PM_2.5_ surges to oxidative stress and JAK2/STAT3 dysregulation, with urban cohorts showing stronger responses and higher relative risk of lung cancer mortality.

## Impact

These results underscore the interplay between PM_2.5_-driven oxidative stress and JAK2/STAT3-mediated oncogenic signaling in both pulmonary and systemic compartments, reinforcing the necessity for improved air quality control, molecular-level monitoring, and targeted mitigation strategies in highly exposed populations.

